# Distinct evolutionary patterns of endemic and emerging parvoviruses, and the origin of a new pandemic virus

**DOI:** 10.1101/2025.07.14.664739

**Authors:** Robert A. López-Astacio, Brian R. Wasik, Hyunwook Lee, Ian E. H. Voorhees, Wendy S. Weichert, Oluwafemi F. Adu, Laura B. Goodman, Susan L. Hafenstein, Uwe Truyen, Colin R. Parrish

## Abstract

Emergence of epidemic viruses in new hosts threatens both human and animal populations, and often involves virus evolution to overcome barriers that normally prevent efficient infection and spread in that host. After transfer the separated viruses will evolve in parallel as they spread within the original and new hosts. Here we examine the details of a virus involved in such a host-jumping event, where we define the natural evolution of feline panleukopenia virus (FPV) over 60 years, clarify the origins of the new pandemic canine parvovirus (CPV) that arose in the 1970s, and compare the separate evolution of those viruses over 47 years in cats or dogs. Several live-attenuated FPV vaccine viruses originated from early 1960s isolates or were a recombinant of an early virus, and many sequences in databases proved to be vaccine-derived. The sequences of wild viruses showed that FPV-like strains evolved at ∼25% the rate seen for CPV in dogs, and the higher rate of CPV evolution was consistent since 1979 when a genetic variant became widespread. The common ancestor of the CPV lineage was related to FPVs from Europe, and contained several unique host-adaptive capsid changes associated with canine transferrin receptor type-1 binding. Although the FPV vaccine strains are around 60 years old, little selection for antigenic variation was observed. The distinct evolutionary patterns of these closely related viruses circulating for decades in different hosts emphasizes the complex evolution associated with viral epidemic emergence and spread in endemic and new hosts.

**SIGNIFICANCE STATEMENT:** Comparing the evolution of a virus in its reservoir host with that seen in a new host will reveal the special circumstances that allow epidemic emergence. A feline parvovirus (FPV) jumping to dogs in the mid-1970s formed canine parvovirus (CPV), which has circulated world-wide until today. The evolutionary rate of FPV in its original hosts was much lower than that of CPV in dogs, and the mutational patterns seen in the different hosts were also distinct. Early CPV isolates differed from the ancestral FPV clade in several key host range mutations. These results highlight the complex biology associated with epidemic emergence, including host-specific rates of lineage evolution and complex origins of host-adaptive mutations. (113/120)

## INTRODUCTION

Control of new viral diseases requires a clear understanding of their evolutionary dynamics, particularly those associated with host shifting allowing epidemic emergence, or of changes in antigenic structures that allow evasion of prior host immunity (1, 2). While viruses that emerge to cause epidemics or pandemics in new hosts are rare, they represent significant threats to human and animal health (3). Our knowledge of how such viruses overcome the many biological, epidemiological and ecological barriers that usually prevent such emergences is incomplete (4–6). In particular we often lack detailed information about the ancestral virus, its sequence or properties, evolutionary patterns, or its transmission in the original reservoir hosts (7). Directly comparing the evolution and biology of viruses in both their original and new hosts can often reveal key details about how such emergence events have occurred, and therefore provide us with the tools to better understand how to forestall similar outbreaks in the future (8, 9).

The emergence of canine parvovirus to cause a pandemic of dogs is a now well-established model of virus emergence where host shift leading to a pandemic (5). The canine parvovirus (CPV) has been most recently classified as *Protoparvovirus carnivoran 1* (10), and its emergence from the feline panleukopenia virus (FPV) as a new virus in dogs occurred around 50 years ago (11). Cerebellar disease in kittens similar to that caused by FPV was reported around 1887 (12), and FPV-like enteric diseases in cats were first reported in the 1920s (13). Enteric diseases in cats and raccoons were widely observed in the 1930s and 1940s, and also experimentally reproduced in cats (14, 15) (reviewed in (16)). In the late 1940s and early 1950s, disease among farmed mink were associated with an FPV-like parvovirus that was named mink enteritis virus (MEV) (17, 18).

The most common diseases are due to FPV replicating in the rapidly dividing cells of the intestinal crypts and bone marrow of cats, resulting in enteritis in kittens older than about 6 weeks, which was often accompanied by panleukopenia (19, 20), and the virus is shed in the feces at high titers (21). Infection of neonatal kittens may result in infection of cells in the cerebellum, resulting in cerebellar hypoplasia and ataxia (22, 23).

FPV was first isolated in tissue culture in the early 1960s in the United Kingdom, and the sample was from a captive snow leopard that likely acquired the virus from a domestic cat (24). That virus or other early 1960s isolates were used to prepare attenuated viruses by extended passage in tissue culture (25). Since the 1970s attenuated FPV vaccine viruses have been included in the core vaccines recommended for all kittens, along with feline herpesvirus (rhinotracheitis) and feline calicivirus (26).

In the mid-1970s a variant of FPV emerged in dogs that was named CPV type-2 (CPV-2) to distinguish it from the previously described minute virus of canines (MVC; now *Bocaparvovirus carnivoran 1*). Although first reported in dogs in many regions of the world during 1978 (27–30), it appears that the CPV-2 strain was circulating in Western Europe for a few years before emerging globally, since anti-CPV antibodies were detected in sera collected from dogs in Greece in 1974 and in Belgium and the Netherlands in 1976 and 1977 (31–33). In other regions of the world (the USA, Japan, and Australia) the first CPV-antibody positive sera were collected in early- to mid-1978 (34–36). Significant strain variation occurred between 1979 and 1981, when the original CPV-2 strain was replaced globally by a genetic variant (CPV-2a) which contained several mutations compared to CPV-2 (37, 38). While the original CPV-2 strain was not able to infect cats, the CPV-2a strain had gained the feline host range (39–41). However, most infections of cats have continued to be due to FPV, so that virus has continued to circulate and evolve in cats in parallel with the CPV strains in dogs (42). As well as cats and dogs, both FPV and CPV strains can naturally infect and cause disease in a variety of other related hosts, which are mainly members of the Order Carnivora (43).

The parvovirus capsid is a 26 nm diameter T=1 icosahedron assembled from 60 copies of a combination of the overlapping VP1 and VP2 proteins, which both are included in the capsid structure so as to form the same small, exposed surface (44, 45). That exposed structure recognizes the host transferrin receptor type-1 (TfR), antibodies, as well as the sialic acid N-glycolylneuraminic acid (Neu5Gc) (46–51). The capsid packages a linear single-stranded DNA genome (ssDNA) of about 5,120 bases which contains two large open reading frames that encoded the non-structural (NS) and the capsid or viral proteins (VP). Alternative splicing produces NS1 and NS2, as well as VP1 and VP2, while proteolysis near the N-terminus of VP2 produces VP3 (52). The virus is transmitted through a fecal-oral route, and the virion is robust and can survive in the environment for weeks, so that viruses may be moved long distances on contaminated fomites (21).

Infection of host cells requires binding to the transferrin receptor type-1 (TfR) (53, 54). TfR is a homo-dimeric type-2 membrane glycoprotein and its most important function for the host is to bind to iron-loaded transferrin (Tf) and transport that into the cell by clathrin-mediated endocytosis (55). The TfR also binds other host proteins, including ferritin and the homeostatic iron regulator protein (also known as High FE2+ (HFE)), in addition to being a receptor for other known pathogens (56). The capsids of CPV and FPV bind to the TfR through a small region on the TfR apical domain, and that interaction has been defined structurally by cryo-electron microscopy (cryoEM) (57, 58) and functionally by mutational analysis of the capsid and receptor (59, 60). The canine host range of CPV-2 (and CPV-2a) is associated with a small number of changes in or near the surface of the viral capsid, which allow the virus to bind the canine TfR and to infect dog cells and dogs (61, 62). Some of the capsid changes between FPV and CPV also altered antibody binding epitopes (46, 63–65), or modified the capsid interactions with the sialic acids present in cats, but not dogs (51, 66). FPV capsids do not bind the canine TfR or infect canine cells, and that is in large-part due to the presence of a fourth N-glycosylation of that receptor apical domain that falls within the capsid binding site (67). The mutations in CPV controlling canine host range allow that capsid to bind to the glycosylated canine TfR - at relatively low affinity - but sufficient to allow that virus to enter and infect dog cells and dogs (63).

The parvovirus is immunogenic in both dogs and cats, and after infection the host rapidly produces protective antibodies against the capsid (68–70). Maternal antibodies transferred in the colostrum protect newborn animals for the first months of life, but also block the infection by attenuated vaccines interfering with vaccine efficacy (71, 72). Analysis of monoclonal and polyclonal anti-parvovirus antibody binding specificities has revealed two dominant antigenic structures on the capsid surface (45–48, 73), and antigenic variation of the capsid may block the binding of individual antibodies (74). However, that antigenic variation has not clearly resulted in functional immune escape of the virus in nature (69, 70, 75–78). In addition, many capsid mutations alter both antibody and TfR binding (47, 67, 79–82), so that antigenicity and host range are coupled targets of selection in parvoviruses.

Previous studies examining the evolutionary patterns of FPV from cats or related hosts mostly examined the VP2 genes, and did not systematically identify the vaccine strains in the datasets (83–89). Among recent studies, several have shown the global distribution of FPV strains and the emergence of a small number of mutations that have spread widely, including several studies identifying a VP2 residue 91 Ala to Ser substitution (90–94). We have recently defined the evolution of the full genomes from natural infections of CPV in detail (95), and here we add a cross-species analysis by similarly defining the evolution of FPV over the past 60 years. We identify the origins of several live-attenuated vaccines in current use, reveal the closest relatives of the CPV-2 lineage among the FPV-like viruses, and describe the different rates and patterns of evolution in the two viral lineages during their decades of parallel spread in cats and dogs.

## MATERIALS AND METHODS

### FPV sample collection

Seventeen FPV-containing samples from different years back to 1964 were obtained from either naturally infected original host tissues or cell cultured isolates; those samples had been either stored in our laboratory or obtained for this study from other collections. The newly sequenced samples are described in **Table 1** (and denoted in **Figure 1** with black triangles), along with other FPV full-genome sequences obtained from the GenBank NCBI database.

**Figure 1.**
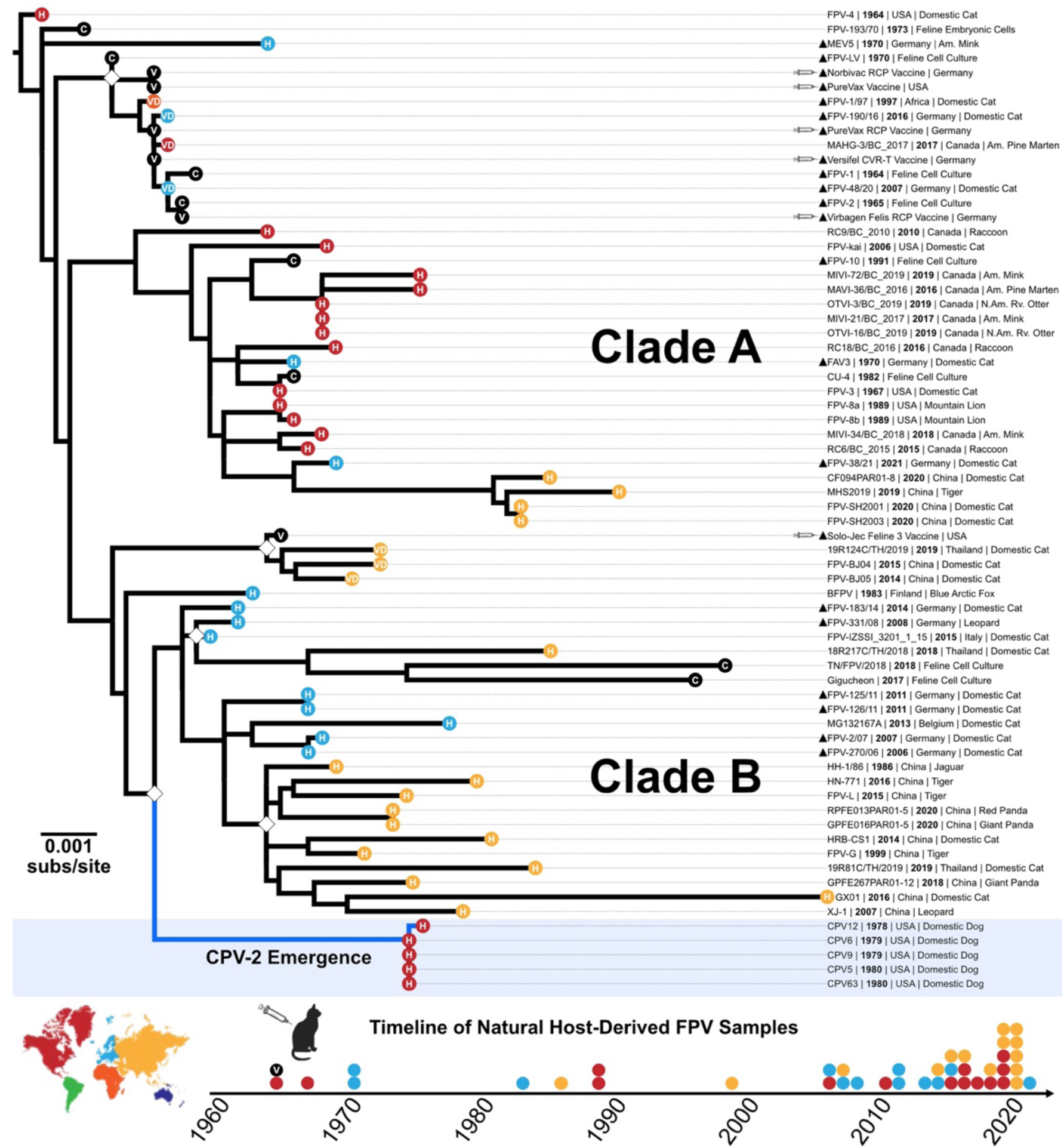
Evolutionary relationships among FPV full-genome sequences. ML tree of full-length FPV genomes (n=63), in addition to an early CPV-2 emergent clade (n=5). Newly isolated sequences are denoted with a triangle. Bootstrap support values of >90% are indicated with a white diamond at the nodes, scale represents nucleotide divergence. Phylogeny tips are denoted with shapes colored by geographic origin (continent) and information of sequence source (C=cell culture-adapted virus; H=host-derived virus, and V=vaccine-derived virus). A timeline of samples (below; colored by country as in the tree) shows the temporal and geographical origins of natural host-derived sequences analyzed.

**Table 1.** Metadata of full-length FPV sequences, including: sample ID, collection year, geographic origin, sample source/host, and GenBank accession. New generate sequences for this study are marked with a triangle (▴).

| FPV VACCINES (n=6) |  |  |  |  |
| --- | --- | --- | --- | --- |
| Sample ID | Collection Year | Country | Source | GenBank Accession |
| ▲ PureVax |  | USA | Live-attenuated vaccine |  |
| ▲ Solo-Jec Feline 3 |  | USA | Live-attenuated vaccine |  |
| ▲ Virbagen Felis RCP |  | Germany | Live-attenuated vaccine |  |
| ▲ Versifel CVR-T |  | Germany | Live-attenuated vaccine |  |
| ▲ PureVax RCP |  | Germany | Live-attenuated vaccine |  |
| ▲ Nobivac RCP |  | Germany | Live-attenuated vaccine |  |
| CELL CULTURE FPV (n=8) |  |  |  |  |
| Sample ID | Collection Year | Country | Source | GenBank Accession |
| ▲ FPV-1 | 1964 | USA | Feline kidney cell culture |  |
| ▲ FPV-2 | 1965 | USA | Feline cell culture |  |
| ▲ FPV-LV | 1970 | Germany | Feline cell culture |  |
| FPV-193/70 | 1973 | Australia | Feline embryo cell culture | X55115 |
| CU-4 | 1982 | USA | Feline kidney cell culture | M38246 |
| ▲ FPV-10 | 1991 | USA | Feline kidney cell culture |  |
| Gigucheon | 2017 | South Korea | Feline kidney cell culture | MN400978 |
| TN/FPV/2018 | 2018 | India | Feline cell culture | MH559110 |
| VACCINE-DERIVED FPV (n=7) |  |  |  |  |
| Sample ID | Collection Year | Country | Host | GenBank Accession |
| ▲ FPV-1/97 | 1997 | Africa | Domestic Cat |  |
| ▲ FPV-48/20 | 2007 | Germany | Domestic Cat |  |
| FPV-BJ05 | 2014 | China | Domestic Cat | MH165482 |
| FPV-BJ04 | 2015 | China | Domestic Cat | MH165481 |
| ▲ FPV-190/16 | 2016 | Germany | Domestic Cat |  |
| MAHG-3/BC_2017 | 2017 | Canada | American Pine Marten | MN862745 |
| 19R124C/TH/2019 | 2019 | Thailand | Domestic Cat | MN127781 |
| NATURAL HOST-DERIVED FPV (n=40) |  |  |  |  |
| Sample ID | Collection Year | Country | Host | GenBank Accession |
| FPV-4 | 1964 | USA | Domestic Cat | EU659112 |
| FPV-3 | 1967 | USA | Domestic Cat | EU659111 |
| ▲ MEV5 | 1970 | Germany | American Mink |  |
| ▲ FAV3 | 1970 | Germany | Domestic Cat |  |
| BFPV | 1983 | Finland | Blue Arctic Fox | MN451652 |
| HH-1/86 | 1986 | China | Jaguar | KX900570 |
| FPV-8a | 1989 | USA | Mountain Lion | EU659113 |
| FPV-8b | 1989 | USA | Mountain Lion | EU659114 |
| FPV-G | 1999 | China | Tiger | MG764510 |
| FPV-kai | 2006 | USA | Domestic Cat | EU659115 |
| ▲ FPV-270/06 | 2006 | Germany | Domestic Cat |  |
| ▲ FPV-2/07 | 2007 | Germany | Domestic Cat |  |
| XJ-1 | 2007 | China | Leopard | EF988660 |
| ▲ FPV-331/08 | 2008 | Germany | Leopard |  |
| RC9/BC_2010 | 2010 | Canada | Raccoon | MF069446 |
| ▲ FPV-125/11 | 2011 | Germany | Domestic Cat |  |
| ▲ FPV-126/11 | 2011 | Germany | Domestic Cat |  |
| MG132167A | 2013 | Belgium | Domestic Cat | KP769859 |
| ▲ FPV-183/14 | 2014 | Germany | Domestic Cat |  |
| HRB-CS1 | 2014 | China | Domestic Cat | KP280068 |
| RC6/BC_2015 | 2015 | Canada | Raccoon | MF069445 |
| FPV_IZSSI_3201_1_15 | 2015 | Italy | Domestic Cat | KX434461 |
| FPV-L | 2015 | China | Lion | MG764511 |
| MAVI-36/BC_2016 | 2016 | Canada | American Pine Marten | MN862744 |
| RC18/BC_2016 | 2016 | Canada | Raccoon | MF069447 |
| HN-ZZ1 | 2016 | China | Tiger | KX685354 |
| MIVI-21/BC_2017 | 2017 | Canada | American Mink | MN862746 |
| MIVI-34/BC_2018 | 2018 | Canada | American Mink | MN862743 |
| GPFE267PAR01-12 | 2018 | China | Giant Panda | MZ357122 |
| OTVI-16/BC_2019 | 2019 | Canada | North American River Otter | MN862748 |
| OTVI-3/BC_2019 | 2019 | Canada | North American River Otter | MN862749 |
| MIVI-72/BC_2019 | 2019 | Canada | American Mink | MN862747 |
| MHS2019 | 2019 | China | Tiger | MN908257 |
| 19R81C/TH/2019 | 2019 | Thailand | Domestic Cat | MN127780 |
| CF094PAR01-08 | 2020 | China | Domestic Cat | MZ357120 |
| FPV-SH2003 | 2020 | China | Domestic Cat | MW811187 |
| FPV-SH2001 | 2020 | China | Domestic Cat | MW650831 |
| RPFE013PAR01-05 | 2020 | China | Red Panda | MZ357119 |
| GPFE016PAR01-05 | 2020 | China | Giant Panda | MW331496 |
| ▲ FPV-38/21 | 2021 | Germany | Domestic Cat |  |

We also obtained and sequenced six commercially available live-attenuated FPV vaccines were obtained and analyzed. PureVax and Solo-Jec Feline 3 (Boehringer Ingelheim), are available in the USA. Virbagen Felis RCP (Virbac), Versifel CVR-T (Zoetis), PureVax RCP (Boehringer Ingelheim), and Nobivac RCP (Intervet International BV).

### FPV genome amplification, DNA library preparation, and sequencing

Total DNA was isolated from viral samples using the E.Z.N.A. Tissue DNA Kit using the manufacturer’s instructions (cat. no D3396-01) and quantified via Qubit 4 Fluorometer (Thermo Fisher Scientific). The DNA was used in polymerase chain reaction (PCR) using the Q5 High Fidelity DNA Polymerase (New England BioLabs) under the conditions specified by the manufacturer. We used 30 cycles to amplify the nearly complete FPV genome as two overlapping segments using the following primers: 5’-CCGTTACTGACATTCGCTTCTTG-3’ and 5’-GAACTGCTCCATCACTCATTG-3’ (for NS1, ∼2.5 kb amplicon), and 5’-CATCCATCAACATCAAGACCAAC-3’ and 5’-CTTAACATATTCTAAGGGCAAACCAACCAA-3’ (for VP2, ∼2.5 kb amplicon). This reaction covered ∼85% of the genome, including the NS1 and VP1 reading frames, but lacked the 5’- and 3’-terminal untranslated regions (UTRs). Each PCR product was purified separately with 0.45 volume AMPure XP beads (Beckman Coulter), quantified via-Qubit 4 Fluorometer, and 0.5ng of each fragment was pooled together as input DNA (1ng DNA total) to construct barcoded sequencing libraries with the Nextera XT DNA Library Preparation Kit (Illumina). Libraries were multiplexed and sequences were determined using Miseq 2×250 Illumina sequencing.

All sequencing reads were deposited at the NCBI Sequence Read Archive (SRA) under BioProject PRJNA1288508.

### Illumina read processing, analysis, and bioinformatics

Raw sequencing reads were trimmed using BBDuk (https://jgi.doe.gov/data-and-tools/software-tools/bbtools/bb-tools-user-guide/bbduk-guide/) to remove adaptors, PCR primers, and low-quality regions from reads. Reads were merged and mapped to a FPV reference sequence (consensus accession no. M38246) using Geneious Prime. Reads were error-corrected and normalized to 5000-fold coverage per site using BBNorm (https://jgi.doe.gov/data-and-tools/software-tools/bbtools/bb-tools-user-guide/bbnorm-guide/). An average of ∼5,000-fold coverage per site was obtained for a majority of the genome. A short region (∼150 nucleotides in length) flanked by a poly(A)- and G-rich sequence region near the start of the VP2 gene and the distal portions of the genome (close to the UTRs) frequently failed to properly assemble. Those low coverage regions (LCR) did not affect consensus calls but would affect intra-host measurements of mutations frequencies.

Consensus genome assemblies were deposited to GenBank, with accession numbers found in Table 1.

### FPV full-genome and VP2 phylogenetic analyses

We determined the evolutionary relationships among the full-genome consensus sequences (n=63), including 22 new sequences (6 from FPV vaccines) prepared here, along with 41 FPV sequences from GenBank for which isolation location, date, and host information were available (**Table 1**). Sequences were aligned using the MUSCLE alignment method (96). Maximum likelihood (ML) phylogenetic analysis was performed using PhyML (97) or IQ-TREE (98), employing a general time-reversible (GTR) substitution model, gamma-distributed (Γ) rate variation among sites, and bootstrap resampling (1,000 replications). As there is no closely related virus outside the FPV (and CPV) collection, trees were either rooted to the earliest FPV sample available (FPV-4, 1964, accession no. EU659112) or were mid-point rooted. The most closely related FPV sequences to the early CPV were identified by including sequences of 1978 and 1979 CPV-2 isolates in the phylogenetic analysis.

Sequences of highly cell cultured viruses and identified vaccine virus genomes from the dataset were removed to allow analysis of the natural infection-derived sequences (n=40) as both full-length genomes, and as NS1 and VP2 sequences separately. We also analyzed a larger dataset of FPV VP2 sequences from the GenBank NCBI database (n=183). Comparison with the CPV natural genomes was done using a dataset of 212 sequences examined in a previous study of that virus (95).

### Mutation and selection of FPV and comparison with CPV

Mutations in the FPV genome found in different categories of samples (wildtype natural infections, vaccine, cell culture) were annotated using the earliest FPV full-genome sequence (FPV-4) as reference. Determining ‘widespread’ polymorphisms in wildtype natural infections was set as a cutoff of ≥10% of samples (at least 4 of 40). The rates of evolution of natural FPV (n=40) and CPV (n=212) genomes was determined using root-to-tip genetic distance by regression of each ML tree by date (year) in the TempEst v.1.5.3 software (99), and also by calculating the divergence rates using a Bayesian Markov chain Monte Carlo (MCMC) approach run in BEAST v.1.10.4 (100). MCMC sampling was performed with a relaxed clock, a generalized time-reversable (GTR) substitution model with gamma distribution in four categories (Γ4), at a minimum of 100 million runs, with duplicates combined in Log Combiner v1.10.4. We confirmed statistical convergence with an effective sample size [ESS] ≥200 (consistent traces, removed burn-in at 10 per cent) in Tracer v1.7.2 (101). Parameter estimation given as Highest Posterior Density (HPD) interval.

The relative numbers of nonsynonymous (dN) and synonymous (dS) nucleotide substitutions per site in NS1 and VP2 ORFs of FPV and CPV natural isolates were calculated using the SLAC (**S**ingle-**L**ikelihood **A**ncestor **C**ounting) method available in the Datamonkey package (https://datamonkey.org/, (102)). A 95% Confidence Interval (CI) was calculated for the mean dN/dSratio and represented as error bars.

### Structural analysis of FPV capsid mutations and receptor or antibody binding sites

The atomic model of the canine TfR was predicted by using Alphafold3 (103) and fitted into the cryoEM density of the black-backed jackal (bbj) TfR in complex with the CPV-2 capsid (EMDB ID: 20002)(PDB ID: 6OAS) (58). The N-glycan moieties [(Hex)2(HexNAc)1(NeuAc)1(Man)3(GlcNAc)2] on the canine TfR residue N384 was modeled into the TfR-CPV complex structure by using CHARMM-GUI Glycan Modeler (104). The effects of viral changes on virus:ligand interactions were examined in the context of cryo-EM structures reconstructed with Chimera X (105). Those included the bbjTfR in complex with the CPV-2 capsid (PDB ID: 6OAS) (58) and the structures of canine antibody Fab-capsid complexes (PDB IDs: 9E7W and 9E60) (106, 107).

### Statistical analyses

Data was analyzed using GraphPad Prism (GraphPad Software, Inc., La Jolla, CA).

## RESULTS

### Sixty years of FPV evolution and identification of vaccine viruses

We employed a phylogenetic analysis of full-length FPV genomes to reconstruct the evolutionary history of that virus, and to define the relationship between vaccine viruses and other isolates. We determined a number of new historic and contemporary FPV sequences, as well as six live-attenuated vaccines widely used in the USA and Western Europe (**Table 1**). We generated a maximum-likelihood (ML) tree of full-length FPV genomes (total n=66), along with three CPV-2 genomes (**Fig. 1**). Most nodes had limited bootstrap support within the phylogeny, reflecting the fact that the sequences share >98.5% nucleotide identity overall. The majority of viruses in our dataset were isolated from domestic cats, but some were collected from other susceptible hosts among the Order Carnivora, including mink, raccoons, pine marten, river otter, pandas, and several large predatory felids.

Most vaccine virus sequences clustered with each other and with other early isolates (24, 25). One vaccine virus (Solo-Jec Feline 3), which grouped with more recent FPV isolates in the full genome phylogeny (**Fig. 1**), contained a VP2 gene closely related to other vaccines and early 1960s isolates (**Fig. 2A**) but had a distinct NS1 sequence, suggesting it is a recombinant virus. A number of sequences that had been reported as wildtype FPV strains clustered with the vaccine viral sequences and were most likely due to inadvertent recovery of vaccine viruses or their DNA (**Fig. 1**).

**Figure 2.**
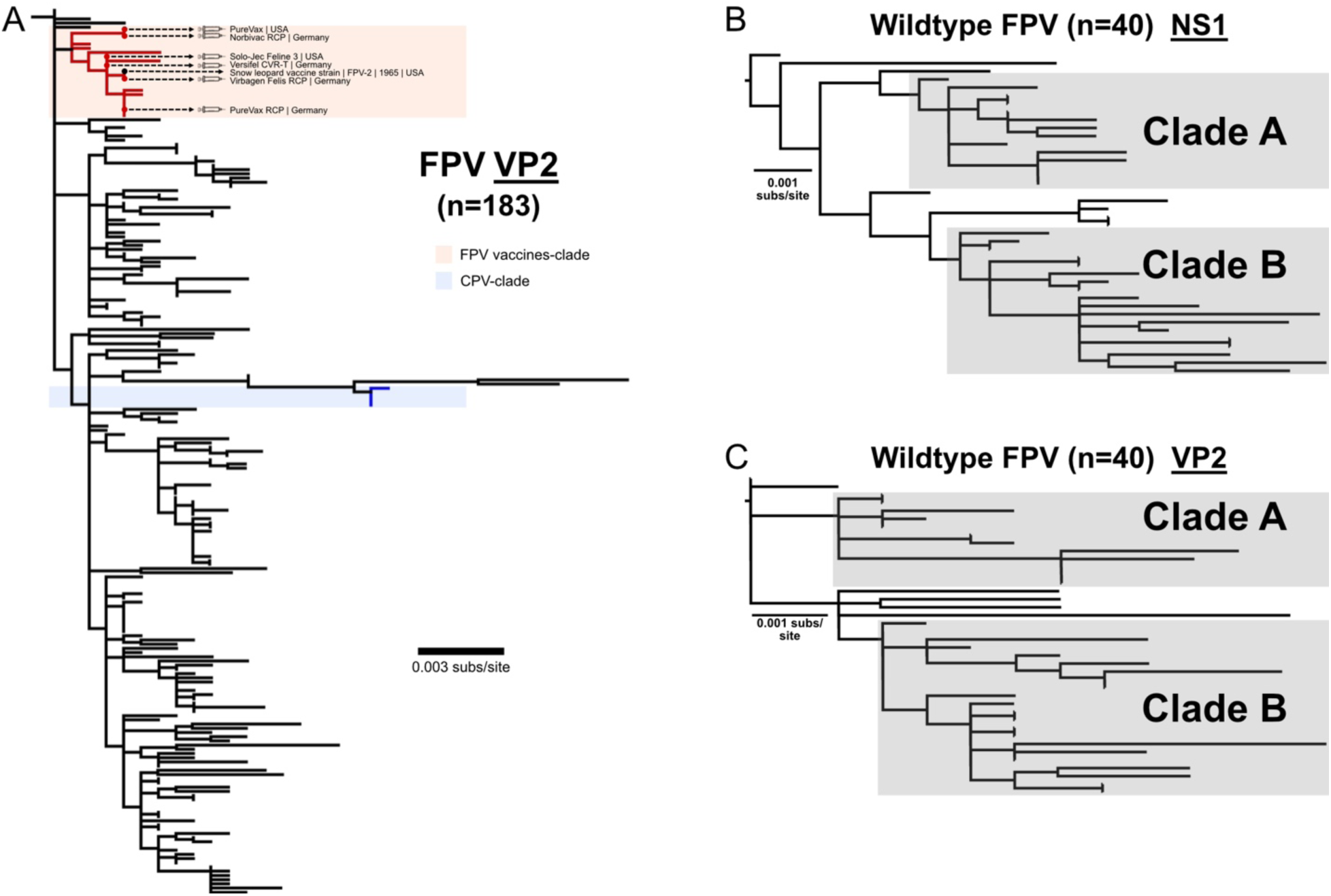
FPV Vaccines have similar VP sequences and FPV natural sequences show convergent organization across genome and absence of natural recombination. (A) ML tree of expanded FPV VP2 sequences (n=183) show close relation of all vaccines in capsid sequence. ML trees of FPV natural sequences (n=40) were generated for both NS1 (B) and VP2 (C) ORFs. Tree structure and clades match that seen with full-length genomes (Fig. 1).

After removing vaccine-derived sequences, as well as those of some highly cell culture passaged viruses, a final collection of 40 FPV wildtype viruses were analyzed as complete genomes. We also separately analyzed the NS1 and VP2 open reading frames, and those showed no obvious genomic recombination lineages among the set of wildtype FPVs (**Fig. 2BC**). The natural sequences fell into two distinct clades (Clade A and B) (**Figs. 1 & 2**). While some FPV isolates from different regions of the world were found in each clade, North American isolates tended to fall within Clade A and European-Asian hemisphere viruses in Clade B.

### The branch leading to the CPV and canine adaptation

The branch leading to CPV-2 fell within the Clade B sequences, and were most closely related to FPVs from Europe collected at various times after the emergence of CPV (**Fig. 1**) (37). That FPV-CPV branch included twelve non-synonymous and ten synonymous substitutions across the genome (**Table 2**). Nine of the non-synonymous substitutions fell within the capsid VP2 open reading frame (**Fig. 3A**). Differences in the VP2 capsid between the ancestral FPV and the earliest CPV-2 sequences included surface residues (80, 93, 232, 323, and 564) adjacent to the 3-fold spike, some of which influence the canine host range (**Fig. 3B**). The CPV-specific mutations (and subsequent CPV-2a replacement) within the capsid protein gene have clear effects on cell binding and infection using the glycosylated canine TfR (**Fig. 3C**) (46, 58, 63, 65, 67).

**Figure 3.**
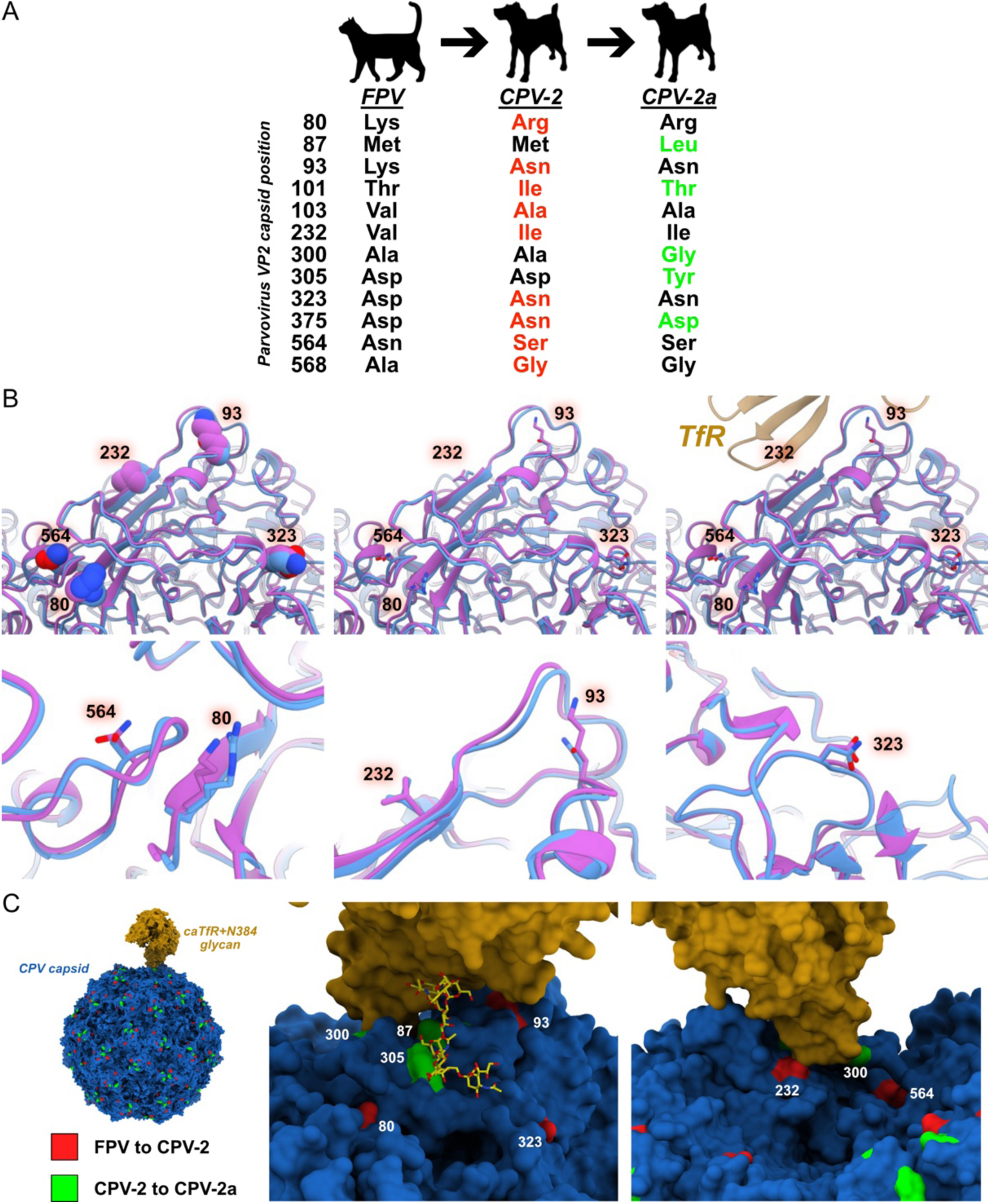
Molecular and structural evolution of parvovirus host shifts. Mutations on branch of CPV-2 emergence. (A) Non-synonymous VP2 mutations fixed during host shifts from likely FPV ancestor to earliest CPV-2 isolates (red) to CPV-2a pandemic (green). (B) Direct comparison of FPV-CPV-2 VP2 side chain changes at 3-fold spike of capsid, and in relation to TfR. (C) Structure of CPV capsid in interaction with canine TfR (including N-glycan structure). Positions in VP2 of mutations arising during shift to CPV-2 (red) and CPV-2a (green) are highlighted on the structure. All mutations present on branch to emergent CPV-2 are listed in Table 2.

**Table 2.** Alignment uncovered variations between consensus FPV lineage to earliest emergent CPV-2 genomes.

| Nucleotide | s/ns | ORF | Amino Acid |
| --- | --- | --- | --- |
| C743T | ns | NS1 | T248I |
| T1224C | s | NS1 | I408 |
| A1479G | s | NS1 | V493 |
| var1620T | s | NS1 | V540 |
| G1926A | ns | NS1 | M152V |
| var2002G | s | NS1 | N/D668 |
| C2110T | ns | VP1 | L4F |
| A2753G | ns | VP2 | K80R |
| A2760G | s | VP2 | V82 |
| A2793C | ns | VP2 | K93N |
| C2816T | ns | VP2 | T101I |
| T2822C | ns | VP2 | V103A |
| G3208A | ns | VP2 | V232I |
| T3213C | s | VP2 | Y233 |
| G3481A | ns | VP2 | D323N |
| A3552G | s | VP2 | E346 |
| A3555G | s | VP2 | A347 |
| G3637A | ns | VP2 | D375N |
| T3681C | s | VP2 | T389 |
| A4137C | s | VP2 | A541 |
| A4205G | ns | VP2 | N564S |
| C4217G | ns | VP2 | A568G |

### Distinct evolutionary patterns of FPV and CPV - and a possible recent common ancestal virus

Examining the 47 years of parallel spread in cats and dogs since 1978 showed significant differences in the rates of evolution of the virus in the original versus new host. Substitution rates of natural isolates of FPV and CPV (n=40 and n=212, respectively) were estimated using both root-to-tip divergence (**Fig. 4A**) and a Bayesian Coalescence Model (**Fig. 4B**). Those approaches showed FPV variation rates of 5.78 x10^-5^ and 5.11 x10^-5^ [HPD 2.03 x10^-5^-7.369 x10^-5^] substitutions/site/year, and CPV rates of 2.181 x10^-4^ and 2.184 x10^-4^ [HPD 1.855 x10^-4^-2.532 x10^-4^] substitutions/site/year. The CPV divergence rate was therefore 3 or 4 times that seen for FPV (**Fig. 4**). This analysis and back-extrapolating the divergence of the FPV lineage also suggested that all of these viruses share a common ancestor that existed around 1890 (HPD 1800-1940).

**Figure 4.**
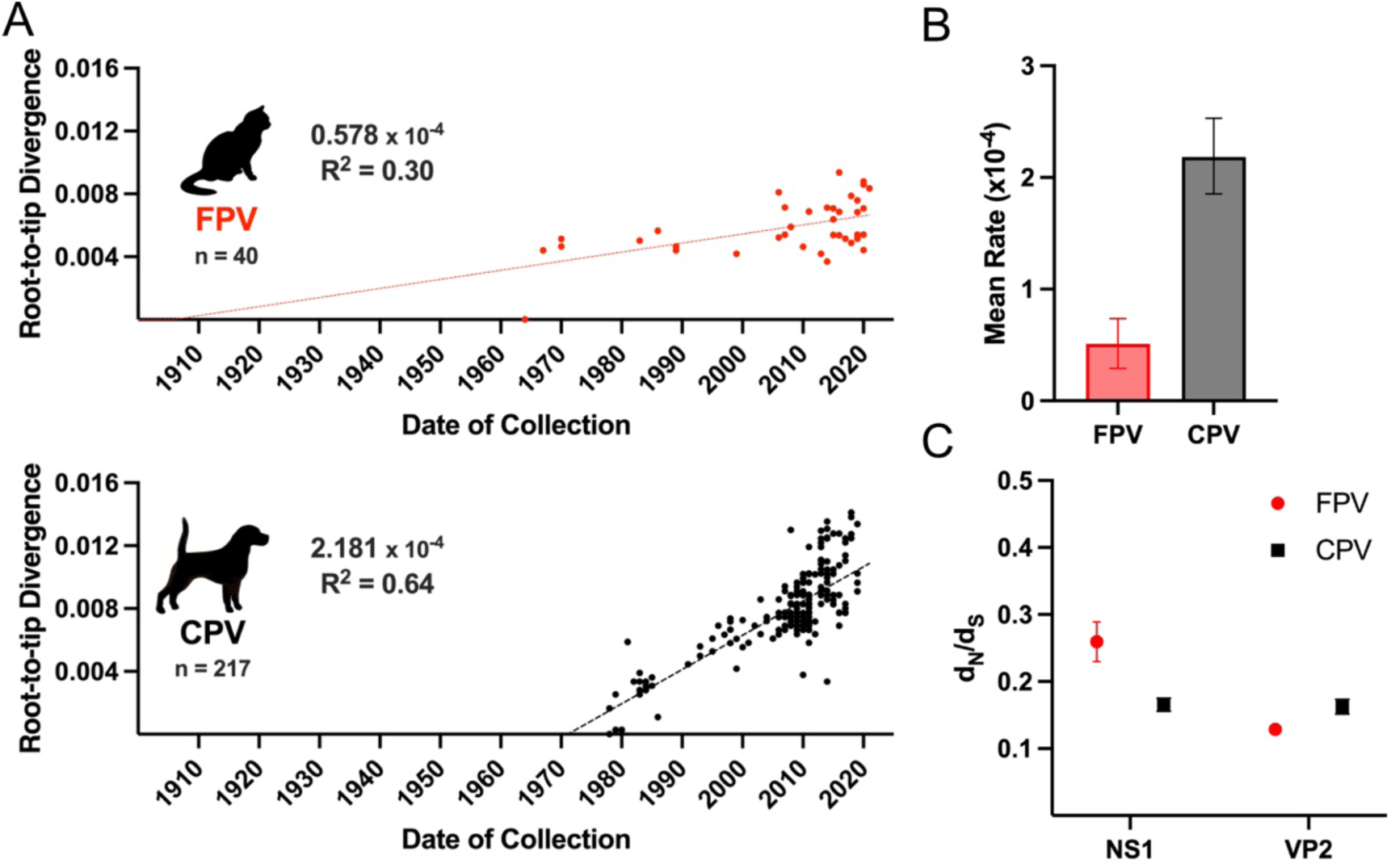
FPV and CPV diverge in evolutionary rates in each host. (A) Divergence rates of FPV (n=40) and CPV (n=217) among recent natural sequences were generated from ML trees analyzed for root-to-tip regression using TempEst. (B) Substitution rates were calculated following a Coalescence Model analysis of natural FPV and CPV sequences. (C) Calculated dN/dS ratios of respective gene ORFs among natural FPV and CPV sequences (Mean +/- 95%CI).

The distinct evolutionary rates of FPV and CPV lineages appeared relatively consistent throughout the periods analyzed, and no significant differences were seen within the CPV evolutionary rates early after emergence in dogs compared to that estimated after the virus had been circulating for 20 years or more. However, a key exception was the single event where five mutations all were fixed together in the ancestor of the CPV-2a lineage (**Fig. 3**).

### Patterns of variation in FPV genomes and comparison to CPV

Comparing the ratios of nonsynonymous to synonymous (dN/dS) mutations among wildtype FPV and CPV sequences over their circulation showed that the ratio for NS1 genes was greater for FPV (0.26 +/-0.11) than seen for CPV (0.17 +/-0.06), while those for the VP2 genes were similar (0.13 +/-0.07 versus 0.16 +/-0.07, respectively) (**Fig. 4C**). There were 45 widespread mutations or polymorphisms among the FPV genomes, including 32 that were synonymous and 13 that were non-synonymous (**Table 3**; **Fig. 5)**. Of the synonymous mutations, 12 fell within the non-structural genes (NS) and 16 within the VP coding region. For the 13 non-synonymous mutations, 6 were found in the NS1 gene, 2 were found to impact both the NS1 and NS2 overlapping reading frames, 2 were in the NS2 gene region alone, 1 was found in the VP2 gene, and 2 were in the small alternatively translated (SAT) alternate open reading frame within the VP2 gene. In contrast to the evolution of CPV over ∼40 years (95), FPV sequences acquired more synonymous and fewer non-synonymous changes (**Fig. 5**). A majority (12 of 23) of the non-synonymous substitutions in CPV are in the VP1/2 capsid gene, compared to just one (of 13 total) in the FPV VP1/2 (**Fig. 5**). The functions of most mutations within the FPV NS1, NS2, and SAT genes are not known, but host specific effects may be possible as these proteins interact with host-derived cellular proteins (108–118).

**Figure 5.**
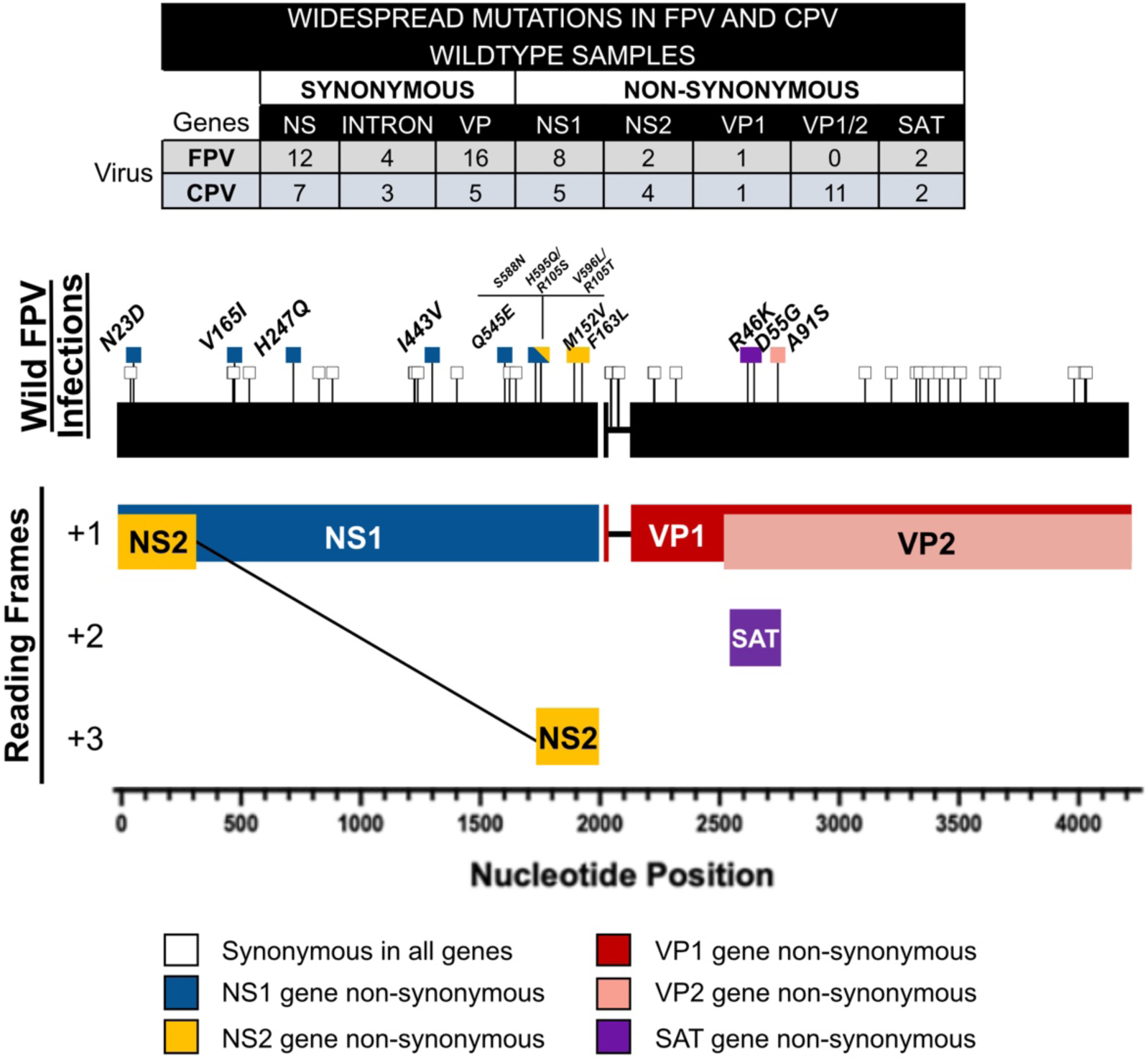
Mutational variation in wildtype FPV genomes. FPV genomes were aligned and categorized by source (wildtype, cell culture, vaccines). Widespread polymorphic variants among wildtype natural infections (≥10%) were determined among FPV (n=40) and CPV (n=217) samples. Mutation counts by type and ORF position are tabled; full listing of widespread variants are found in Table 3. FPV mutations are visualized on the genome by pins (non-synonymous = colored squares, synonymous = white squares). Nucleotide numbering begins at the first coding site of the NS1/2 gene. Amino acid numbering is based on each respective gene translational start site.

**Table 3.** Widespread variants among FPV wildtype natural infection samples (n=40).

| Nucleotide | s/ns | ORF | Amino Acid | % samples |
| --- | --- | --- | --- | --- |
| G54A | s | NS1 | L18 | 25 |
| A67G | ns | NS1 | N23D | 42.5 |
| T486G | s | NS1 | L162 | 12.5 |
| T489C | s | NS1 | C163 | 10 |
| G493A | ns | NS1 | V165I | 15 |
| G555A | s | NS1 | V185 | 10 |
| T741A | ns | NS1 | H247Q | 45 |
| G849T | s | NS1 | V283 | 27.5 |
| C906T | s | NS1 | D302 | 25 |
| G1251A | s | NS1 | V417 | 15 |
| T1254G | s | NS1 | G418 | 32.5 |
| T1266C | s | NS1 | C422 | 12.5 |
| A1327G | ns | NS1 | I433V | 47.5 |
| T1431A | s | NS1 | I477 | 12.5 |
| C1633G | ns | NS1 | Q545E | 57.5 |
| T1653C | s | NS1 | Y551 | 35 |
| A1680G | s | NS1 | E560 | 37.5 |
| G1763A | ns | NS1 | S588N | 15 |
| C1785A | ns | NS1/NS2 | H595Q/R105S | 45 |
| G1786C | ns | NS1/NS2 | V596L/R105T | 10 |
| A1926G | ns | NS2 | M152V | 40 |
| T1959C | ns | NS2 | F163L | 57.5 |
| G2080A | s | intron |  | 37.5 |
| T2083A | s | intron |  | 10 |
| C2110T | s | intron |  | 12.5 |
| T2114C | s | intron |  | 10 |
| T2262C | s | VP1 | Y59 | 47.5 |
| C2265T | s | VP1 | F60 | 52.5 |
| A2355G | s | VP1 | K90 | 47.5 |
| G2658A | ns | SAT | R46K | 12.5 |
| A2685G | ns | SAT | D55G | 45 |
| G2785T | ns | VP2 | A91S | 12.5 |
| A3153G | s | VP2 | Q213 | 12.5 |
| G3264A | s | VP2 | V250 | 10 |
| G3369A | s | VP2 | L285 | 12.5 |
| C3385T | s | VP2 | L291 | 52.5 |
| C3420T | s | VP2 | N302 | 12.5 |
| A3468G | s | VP2 | Q318 | 10 |
| T3504A | s | VP2 | I330 | 12.5 |
| G3555A | s | VP2 | A347 | 55 |
| A3663G | s | VP2 | Q383 | 10 |
| T3699C | s | VP2 | P385 | 32.5 |
| G4035A | s | VP2 | T507 | 65 |
| A4080G | s | VP2 | V522 | 35 |
| T4086C | s | VP2 | Y524 | 62.5 |

FPV samples collected from hosts other than cats showed few specific sequence patterns compared to the feline isolates (83, 86, 119, 120), although a VP2 G299E mutation was seen in giant panda isolates (121, 122).

### Capsid mutations, FPV vaccines and antigenic variation

Live-attenuated FPV vaccines analyzed were all closely related to early FPV isolates, including the snow leopard strain from around 1964 (FPV-2) (**Fig. 1**). The vaccine strains often have unique or shared non-synonymous changes in NS1 and NS2. Two VP2 non-synonymous changes were shared among the majority of vaccine strains, including the V232I change on the 3-fold spike and V562L was near the 2-fold axis, which were both exposed on the capsid surface (**Fig. 6A**). Residue VP2 232 fell within the TfR binding footprint, while VP2 562 was immediately adjacent (**Fig. 6B**). FPV vaccine variant residue 232 is within or close to both canine antibody binding sites (45), while VP2 residue 562 is adjacent to one site (**Figs. 6C-D**).

**Figure 6.**
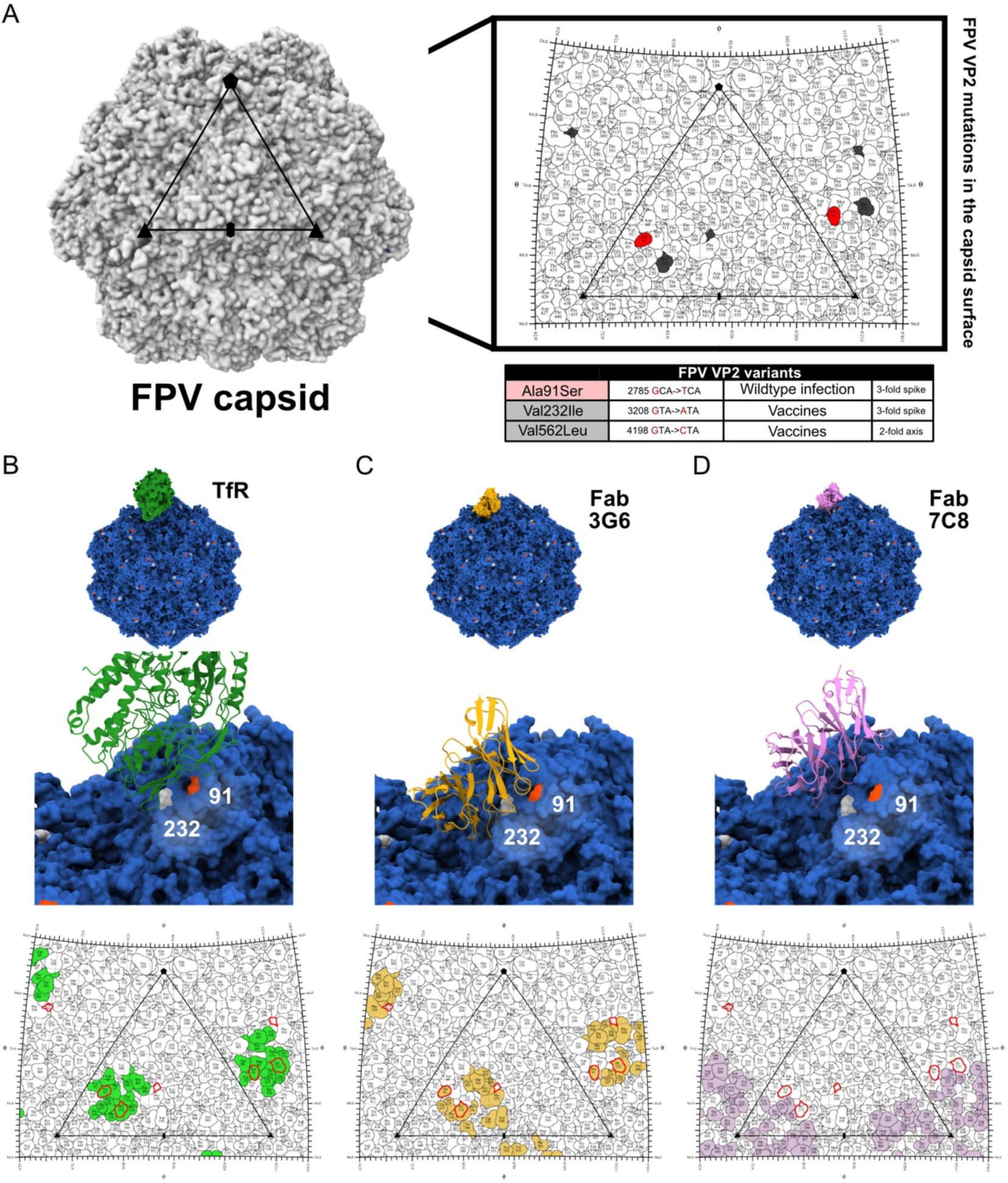
Widespread FPV VP2 and their relative position in the virus capsid surface in context with TfR and antibody footprints. (A) Reconstruction of the FPV capsid, axes of symmetry (oval= twofold axis, triangle= threefold axis, and pentagon=fivefold axes), and a roadmap of a single icosahedral asymmetric unit (PDB ID: 1C8G). The table below the roadmap identifies the widespread variants shared among wildtype infection samples (red) or majority of vaccine strains (grey). (B) Reconstruction of the high-resolution structure of the parvovirus capsid in complex with TfR (green, PDB ID: 6OAS) with corresponding roadmap where variants are outlined in red relative to the TfR footprint (green). (C) Reconstruction of the FPV capsid in complex with a canine-derived monoclonal antibody as Fab: 3G6 (yellow, PDB ID: 9E7W) with corresponding roadmap where variants are outlined in red relative to the Fab3G6 footprint (yellow). (D) Reconstruction of the FPV capsid in complex with a canine-derived monoclonal antibody as Fab: 7C8 (purple, PDB ID: 9E60) with corresponding roadmap where variants are outlined in red relative to the Fab7C8 footprint (purple).

The only consistent variant found among wildtype FPVs was VP2 A91S which is exposed in the 3-fold spike (**Fig. 6A**) within the TfR footprint (**Fig. 6B**) and the antibody binding site of canine antibody Fab2C5 (**Fig. 6C**) and adjacent to antibody Fab7C8 (**Fig. 6D**).

## DISCUSSION

Here we show that FPVs in cats and related carnivore hosts evolved slowly over more than 100 years, forming two clades that are partially geographically structured, with a single branch from one clade giving rise to the ancestor of the CPV lineage. The FPVs in cats or other susceptible hosts accumulated few widely distributed coding changes in their NS1 and VP genes, but the branch leading to CPV included several canine host range–adaptive mutations, highlighting the complex processes required for the virus to overcome the host barrier and to spread widely in its new host. Although there have been many studies describing aspects of FPV and CPV evolution, in this analysis we are able to specifically analyze wildtype FPVs collected from natural infections over 60 years, and to identify the vaccine viruses within this evolutionary context. This detailed understanding of the sequence evolution of FPV can be compared directly to that seen for CPV, as well as to our improved understanding of the structures and functions of FPV and CPV capsids and their interactions with feline, canine and black-backed jackal TfRs and also of host antibodies from infected dogs.

### FPV strains show distinct patterns of variation compared to CPV

Over the 47 years of parallel spread in different hosts the FPV sequences evolved 0.5 x10^-4^ subs/site/year, compared to 2.2 x10^-4^ subs/site/year for CPV (**Fig. 4AB**). The evolutionary rates of CPV sequences between 1979 and 1989 and between 1989 and 2024 were similar indicating that was a steady rate for the virus once it emerged in dogs – with the apparent exception of an early burst of evolution that gave the CPV-2a strain which had 7 new coding changes, allowing that virus to sweep the original CPV-2 to extinction between 1979 and 1981 (37, 38, 64). It is unclear how that group of CPV-2a mutations arose and were selected, but the emergence of partial intermediate viruses in raccoons may have allowed combinations of mutations to arise that would be inviable in dogs (123). Of the wide-spread mutations that arose in the FPV lineage only VP2 V232I was seen among the CPV sequences (95), indicating there were different selections or genetic drift in each host. The overall faster evolution of the CPV lineage may be due to faster genetic drift resulting from different epidemiological properties of virus in dogs, and/or to continuing host or antigenic selection. The rate seen for FPV in this study is similar to that determined for human B19 parvoviruses using combinations of contemporary- and ancient DNA-derived sequences, which were calculated to be 1 to 3 x10^-5^ subs/site/year (124, 125).

### Changes Selected in Alternative Hosts

Cat and dog populations vastly outnumber those of other carnivore hosts for these viruses, and so we expect that most FPV and CPV evolution occurs in those animals. No host-specific variation was seen in FPVs from snow leopards, wild cats, minks, raccoons, or bobcats, while variation of VP2 residue 299 (Ala to Glu) has been reported in FPVs infecting giant pandas (121, 122). In contrast CPVs from non-canine hosts often show variation in capsid sites associated with TfR binding (123, 126), likely due to the capsid structural changes needed to bind and infect using the non-glycosylated TfRs in those hosts (46, 63, 67).

### Sixty-year-old FPV vaccine strains select little antigenic variation

All current FPV live-attenuated vaccines appear to be derived from one or a few 60 year-old isolates, and those are core feline vaccines used in many countries around the world (26). We observed only one or two variations in the residues within the two main epitopes of the FPV capsid epitopes, suggesting significant antigenic selection and immune escape had not occurred in the virus infecting cats. In CPV a small number of antigenic variants have also been observed, but without clear evidence of immune escape occurring (78, 79). While escape from monoclonal antibody selection occurs readily in parvoviruses in *in vitro* studies (74), the host polyclonal antibodies appear to efficiently block infection and circulation of all strains of the virus in its animal hosts (20, 127). In this property the FPV (and CPV) appear similar to other viruses where vaccines have retained their efficacy for decades with little immune evasion of the viruses they are protecting against, such as smallpox, measles, rinderpest, and polio viruses (128–131).

### All FPV (and CPV) strains share a recent common ancestor, and CPV-2 is most closely related to European FPV strains

Phylogenetic and rate analysis of the FPV and CPV sequences suggest that those viruses all share a common ancestor from the late 19^th^ century. This may indicate a novel introduction of an FPV into cats around that time, or the global sweep of a single viral sequence. The first reports of feline ataxia due to cerebellar hypoplasia in two kittens, a characteristic disease caused by FPV, were from 1887 (12), while other FPV-associated diseases in cats and raccoons were reported in the 1920s and 1930s (13–15). However, it is possible that back-extrapolation of the sequences collected since the 1960s does not reflect the longer-term evolution of the viruses, and those have been circulating in a similar form for much longer. This could be resolved by analysis of ancient DNA of viruses in cat mummies or similar samples. Recent sequences of the human B19 parvovirus appeared to have a common ancestor in the 19^th^ century (132), but ancient DNA in human remains up to ∼7000 years old showed that the same strains are much older (124, 125). Sequences of FPVs in ancient cat samples would likely resolve this question.

The CPV-2 strain was recognized when it spread worldwide in 1978, when the first viral sequences were collected (11, 29, 30). However, serological testing showed that the virus circulated in Western Europe for a few years before 1978 (31–33). While the ancestor of the CPV lineage fell within a group of FPV viruses from Europe (**Figs. 1** and **2**), those European FPV isolates were collected between 2006 and 2016, so may be descents of that virus. The true ancestor may still be sought among archival tissues collected from dogs, cats or related hosts in Europe during the early to mid-1970s. This analysis also rules out the possibility that CPV emerged from an FPV vaccine (**Fig. 1**) (133, 134).

### Acquisition of the canine host range

The CPV to FPV branch included nine synonymous and twelve non-synonymous changes – and the latter included the several host adaptive changes, including VP2 residues 93, 103, 323, and 375 (**Fig. 3A**) (46, 135). While a few examples of variation of those sites have been reported (e.g. VP2 93 Lys in a CPV background (88), VP2 93 Asn in an FPV background (89)) those appear to be rare exceptions. Canine host range resulted from the virus adapting the capsid contacts with the canine TfR, including both protein-protein and protein-glycan interactions (53, 61, 67, 136). The canine adaptive mutations each resulted in only small changes in protein structure, mostly of amino acid side-chains and one or two hydrogen bonds, which together allowed them to engage the canine TfR to allow entry and infection (**Fig. 3B and 3C**) (58, 81).

### What does this tell us about viral emergence?

This study confirms that FPV-like viruses have been infecting cats and related hosts for over 100 years and have evolved at a low rate over that time. While dogs would have been consistently exposed to FPV, CPV only emerged once to cause a sustained epidemic. The slow evolution of FPV and long branch to CPV indicates a special process introduced and retained the several canine host range mutations. The ssDNA parvovirus genome is replicated by host cell DNA polymerase during cellular S-phase (most commonly pol Δ), although likely without the error correcting mechanisms that apply during host genome replication (137). In this study polymorphisms were rarely detected within the deep sequenced FPV DNA samples (as was also seen for natural CPV samples (95)). However, one or two base changes were quickly selected when viruses were grown in different host cells, or in the presence of neutralizing antibodies (74, 95). Some canine-adaptive mutations resulted in loss of the feline host range, so would only be selected in dogs or a non-feline host (39). Several mutations together also occurred in the emergence of CPV-2a, and those may have been facilitated by passage through an intermediate host such as raccoons (37, 38).

In many cases it is not possible to directly comparing the properties and evolution of a virus before and after pandemic emergence, since the ancestral virus cannot be analyzed in its original host from the period before emergence. Directly comparing FPV and CPV both reveals the nature of the viruses in their reservoir hosts, as well as the process of successful host-jumping and onward spread.

## ACKNOWLEDGEMENTS

We thank Dr. Edward Holmes (University of Sydney) for creative collaborations related to the evolution of viral emergence. Antibody:capsid structures and molecular analyses were performed with UCSF Chimera X, developed by the Resource for Biocomputing, Visualization, and Informatics at the University of California, San Francisco, with support from National Institutes of Health R01-GM129325 and the Office of Cyber Infrastructure and Computational Biology, National Institute of Allergy, and Infectious Diseases.

## SUPPORT

Supported by National Institutes of Health grants R01-AI092571 and R01-GM080533 to Colin R. Parrish, NIH-Diversity Supplement 3R01AI092571-08W1 to Colin R. Parrish and Robert López-Astacio, and The Daversa Family Scholarship to Robert López-Astacio.

## Notes

### Competing Interest Statement

The authors have declared no competing interest.

